# Transcriptional profiles of functionally distinct HLADR^+^CD38^+^ CD8 T cells subsets from acute febrile dengue patients

**DOI:** 10.1101/2022.09.09.507387

**Authors:** Prabhat Singh, Prashant Bajpai, Deepti Maheshwari, Yadya M Chawla, Kamalvishnu Gottimukkala, Elluri Seetharami Reddy, Keshav Saini, Kaustuv Nayak, Sivaram Gunisetty, Charu Aggarwal, Shweta Jain, Chaitanya, Paras Singla, Manish Soneja, Naveet Wig, Kaja Murali-Krishna, Anmol Chandele

## Abstract

Previous studies showed that a discrete population of the CD8 T cells with HLADR^+^CD38^+^ phenotype expand massively during the acute febrile phase of dengue natural infection. Although about a third of these massively expanding HLADR^+^CD38^+^ CD8 T cells were of CD69^high^ phenotype, only a small fraction of them produced IFNγ upon *in vitro* peptide stimulation. What other cytokines/ chemokines do these peptides stimulated HLADR^+^CD38^+^ CD8 T cells express, what transcriptional profiles distinguish the CD69^+^IFNγ^+^, CD69^+^IFNγ^-^, and CD69^-^IFNγ^-^ subsets, and whether the expansion of the total HLADR^+^CD38^+^ CD8 T cells or the IFNγ producing CD8 T cells differ depending on disease severity remained unclear. This study addresses these knowledge gaps. We find that the CD69^+^IFNγ^+^ subset uniquely expressed key genes involved in protein translation, cellular metabolism, proliferation and dendritic cell cross talk. Both the CD69^+^IFNγ^+^ and CD69^+^IFNγ^-^ subsets had an antigen responsive gene signature with genes involved in cytotoxic effector functions, regulation of T cell receptor signaling, signaling by MAPK, chemotaxis and T cell trafficking to inflamed tissues with the expression being more robust in the IFNγ^+^ CD69^+^ subset. On the other hand, the CD69^-^ IFNγ^-^ subset was biased towards expression of genes that both augment and dampen T cell responses. Lastly, the expansion of total HLADR^+^ CD38^+^ CD8 T cells and also the IFNγ producing HLADR^+^ CD38^+^ CD8 T cells was similar in patients with different grades of disease. Taken together, this study provides valuable insights into the inherent diversity of the effector CD8 T cell response during dengue.

## Introduction

Dengue is a global epidemic resulting in over 100 million clinically significant human cases worldwide each year with India contributing to nearly a third of the global dengue disease burden (1, 2). This mosquito borne acute systemic viral infection can result in clinical disease with symptoms ranging from mild febrile illness (dengue infection without warning signs, DI) to dengue with warning signs (DW) and severe dengue (SD), that can sometimes be life-threatening and fatal (3). Numerous studies conducted over the past several decades suggest that a multitude of host immune factors influenced by the viral-host interface, ranging from an exaggerated host innate and adaptive immune responses and inflammatory cytokines, are associated with immunopathology (4-10). Currently there is no universally licensed vaccine. Numerous vaccines are under research and evaluation and one vaccine CYD-TDV has gone through the most advanced clinical trials is licensed in some countries. This tetravalent dengue CYD-TDV vaccine expressing dengue structural proteins prM and Envelope on a yellow fever vector backbone mainly induces antibody responses since it does not carry dengue nonstructural proteins, which are major targets of the T-cell mediated response. Interestingly, this vaccine has shown demonstrable efficacy in dengue pre-immune individuals while enhancing disease in dengue naïve individuals (11) -thereby further strengthening the view point that, perhaps, vaccines that target CD8 T cells (in addition to antibody responses) may be needed for optimal protection against dengue.

For devising the successful vaccines, it is important to gain a detailed understanding of the human T and B cell responses and their association with protection or immunopathology during natural infection. Towards this, numerous studies revealed that while antibodies are important for protection, they can also enhance infection through a process defined as ‘antibody dependent enhancement’ (ADE) (4, 5, 12). Furthermore, some of the early historical studies hypothesized that cross-reactive memory T cell responses may also contribute to dengue immunopathology via mediating a “cytokine storm” that might directly or indirectly enhance disease. This hypothesis was initially supported by early studies in 1990’s that showed expansion of a CD69 expressing CD8 T cell population peaked around the time when the patients manifested severe disease and hemorrhage (13, 14). However, several recent studies from both murine models and humans raised the possibility that the CD8 T cell responses are probably not drivers of disease severity (15-18). Therefore, there has been a renewed interest towards a detailed characterization of human CD8 T cell responses to dengue natural infection as well as to develop and evaluate vaccines that can induce both antibody and T cell responses. Previous studies from our lab and that of others showed that while there is a massive expansion of HLADR and CD38 expressing CD8 T cell population in dengue patients, only a small portion of these massively expanding CD8 T cells produced IFNγ in response to dengue peptides *ex vivo* (19-21). However, several knowledge gaps remain: What other cytokines/ chemokines do these massively expanding HLADR^+^ CD38^+^ CD8 T cells express upon peptide stimulation? What transcriptional profiles distinguish the CD69^+^IFNγ^+^, CD69^+^IFNγ^-^, and CD69^-^IFNγ^-^ populations among these massively expanding HLADR^+^ CD38^+^ CD8 T cells? Does the expansion of the total HLADR^+^CD38^+^ CD8 T cells or the IFNγ producing CD8 T cells differ depending on disease severity?

This study addresses these questions by RNA seq analysis of the sorted HLADR^+^CD38^+^ CD8 T cells that were stimulated with dengue peptides and sorted as - CD69^-^IFNγ^-^, CD69^+^IFNγ^-^ or CD69^+^IFNγ^+^, and then compared to unstimulated HLADR^+^CD38^+^ CD8 T cells and to HLADR^-^CD38^-^ CD8 T cells that are enriched in naïve CD8 T cells. We found that both CD69^+^IFNγ^-^ and CD69^+^IFNγ^+^ subsets were very similar and were highly enriched in signatures of functional antigen reactive cytotoxic effector T cells with genes like GzmB, GzmA, Prf1, Gnly, transcription factors like Tbx21, Prdm1 and co-stimulatory molecules such as TNFSF9 (4-1-BB), CD27 and ICOS. The CD69^+^IFNγ^+^ subset not only robustly expressed these common antigen reactive cytotoxic effector gene signatures but also preferentially expressed key genes involved in protein translation, cellular metabolism, proliferation and dendritic cell cross talk and several genes of the protein translation machinery that perhaps allowed for an overall heightened expression of these antigen-responsive genes including IFNγ. On the other hand, the bulk of the CD69^-^IFNγ^-^ HLADR^+^CD38^+^ CD8 T cells subset expressed several cytokines and chemokines that are associated with dampening of the T cell response, combined with co-stimulatory molecules and enzymes that augment T cell responses. Lastly, the total HLADR^+^CD38^+^ CD8 T cells and the IFNγ producing HLADR^+^CD38^+^ CD8 T cells were similar in patients with different grades of disease severity strengthening recent studies that have de-linked CD8 T cell responses from dengue disease outcome. Taken together, this study provides valuable insights into the diversity of the effector CD8 T cell response and reveals distinct functional subsets of the massively expanding HLADR^+^CD38^+^ CD8 T cells during natural dengue virus infections.

## Methods

### Human samples

This study uses PBMC and plasma samples obtained from dengue confirmed acute febrile patients that were recruited at the Department of Medicine, All India Institute of Medical Sciences (AIIMS, New Delhi), India during the years 2018-2021. A portion of the blood sample collected for routine clinical tests at the time of enrollment, when available in sufficient quantity, was sent to research laboratory for T cell analysis. The study was approved by institutional ethical boards of the participating institutions. Informed consent was taken from enrolled participants.

### Dengue confirmation

For dengue virus infection confirmation, Dengue NS1 Elisa (J Mitra, IR031096) and Dengue IgM Elisa (Pan Bio, 01PE20) was used in combination. Only those samples from patients who are confirmed for either dengue NS1 and/or IgM and negative for malaria (SD Bioline, 05EK40) and chikungunya (J Mitra, IR061010) were included in this study.

### Disease classification

Disease severity grade was scored at the time of recruitment using the WHO 2009 criteria of Dengue infection without warning signs (DI), Dengue with warning signs (DW) and Severe Dengue (SD) (3).

### Whole blood processing and PBMC / plasma isolation

The Vacutainer CPT tube (Becton Dickenson, Cat# 362761) containing blood sampling was centrifuged at 1500g for 25 minutes without brake at room temperature. After centrifugation, the uppermost layer containing plasma was aspirated and then transferred to cryo-vial tubes and stored at -80°C. The white buffy coat above the gel plug containing the PBMCs was transferred to sterile 15 ml falcon tube and filled with phosphate buffered saline (PBS) (HiMedia #TL1099) and centrifuged at 1800 RPM for 8 min at 4^°^C. After centrifugation the supernatant was discarded, the pellet was re-suspended and washed twice in PBS. After washing, PBMC pellet was re-suspended in red blood cell lysis buffer (HiMedia Cat#R075) and incubated for 2 min to remove any remaining RBCs. The 15 ml tube was then filled with RPMI 1640 media (HiMedia #AL162S) containing 1% fetal bovine serum (FBS) and centrifuged at 1500 RPM for 5 min at 4 C and washed twice. The pellet was re-suspended in 1ml of RPMI media containing 10% FBS and the cells were counted using 0.1% trypan blue (HiMedia #TCL046). The cells were either used immediately or cryopreserved in liquid nitrogen in FBS (Hyclone #SV30014.03) with 10% dimethyl sulfoxide (MP #196055) for later analysis.

### Analytical flow cytometry

PBMCs were washed and surface stained for 30 minutes in ice cold FACS buffer at 4^°^C (1xPBS containing 0.25% bovine serum albumin). Fixable viability dye e-Fluor 780 (Ebioscience, 65086518) was used for excluding dead cells at the time of analysis. For CD8 T cells subsets analysis, surface staining on PBMCs was performed with CD3 (clone UCHTI), CD8 (clone SK1), CD38 (clone H1T2) and HLADR (clone L243) antibodies.

For intracellular staining, cells were fixed with fixation buffer and then permeabilized with Cytofix / Cytoperm (e-bioscience) followed by staining for 60 minutes with antibodies that were diluted in 1X perm buffer (BD, 554723). Flow cytometry data acquisition was performed either on LSR Fortessa X-20 (BD) or FACS Canto II. Data was analyzed using FlowJo software (TreeStar Inc.). CD8 T cells were gated as those that co-expressed CD3 and CD8 and were analyzed of phenotypes and functions.

### Ex vivo stimulation and functional assessment of CD8 T cells

Unless otherwise mentioned, isolated PBMCs were plated at 1×10^6^ cells/well in 96 well U-bottom plates and cultured with pool of overlapping peptides of either entire dengue proteome at a concentration of 1ug/ml of each peptide along with co-stimulation with purified anti-human CD28 (clone L293) and CD49D (clone L25). Unstimulated cells were used as negative control. As a positive control, cells were also stimulated with 1X cell stimulation cocktail containing PMA and ionomycin (Ebioscience, 00-4970-03,). Cells were then cultured for 2 hours at 37°C in 5% CO2 incubator and followed by the addition of protein secretion inhibitor cocktail containing brefeldin A and monensin (GolgiPlug, BD, 555029) for another 4-hours. The cells were then harvested; surface stained with antibody cocktail containing fixable viability dye, CD3, CD8, CD38 and HLA-DR and then fixed and permeabilized using Cytofix/Cytoperm Kit. Cells were then processed for intracellular staining with antibodies against IFNγ (clone 4S.B3), granzyme B (clone GB11) or perforin (clone deltaG9). Cells were acquired on BD Canto II or BD LSRFortessa X-20 and analyzed using FlowJo (TreeStar).

### IFNγ capture assay to identify and sort IFNγ producing CD8 T cells

For identification and sorting of viable IFNγ secreting cells, we used human IFNγ secretion assay detection kit (PE) (Miltenyi Biotec, #130-054-202) as per the manufacturer’s protocol. Briefly, PBMCs were cultured in 12 well flat-bottom plate and stimulated with dengue CD8 megapools (18) that were prepared at a final concentration of 1ug/ml for each peptide. Unstimulated cells were used as controls. Cells were cultured in a tissue culture incubator at 37°C for 3 hours in presence of 5% CO2. The cells were then mixed with IFNγ catch reagent (PE conjugated anti-human IFNγ mouse monoclonal IgG1 conjugated to anti-human CD45 mouse IgG2a). The cells were then allowed to secrete IFNγ for an additional 45 minutes and then IFNγ secreting cells were identified by labeling with PE-conjugated IFNγ detection antibody (anti-human IFNγ mouse IgG1) conjugated to PE).

### Flow cytometric cell sorting for IFNγ producing and non-producing CD8 T cells

After staining the cells with IFNγ capture reagent as described in the section above, the cells were surface stained with fixable viability dye, CD3 (clone UCHT1), CD4 (clone RPA-T4 / OKT-4), CD8 (clone SK1), CD38 (clone H1T2), HLADR (clone L243) and CD69 (clone FN50). Cells were then analyzed in BD FACS Aria Fusion II. After lymphocyte and doublet discrimination, we first gated on CD3^+^, then excluded CD4^+^ cells. After this, the gated CD8^+^CD3^+^ population was analyzed to distinguish peptide stimulated HLADR^+^CD38^+^CD69^+^IFNγ^+^, peptide stimulated HLADR^+^ CD38^+^CD69^+^IFNγ^-^ and peptide stimulated HLADR^+^CD38^+^CD69^-^IFNγ^-^. These CD8 T cell subsets of interest were then sorted in lysis buffer at 4°C using BD FACS aria II cell sorter. As an additional control, we also sorted the total HLADR^+^CD38^+^CD8 T cells and HLADR^-^CD38^-^CD8 T cells that were not stimulated with dengue peptide pool.

### RNA isolation and library preparation for RNA seq

RNA sequencing libraries from the bulk sorted cell subsets were prepared with Illumina compatible SMARTer Stranded Total RNA-Seq Kitv2 - Pico Input Mammalian (TakaraBio, Inc. CA, USA) at Genotypic Technology Pvt. Ltd., Bangalore, India. For each condition approximately 200-500 cells were sorted and collected in 10x lysis buffer of SMART-Seq v4 ultra low input RNA seq kit. From this 8μl was taken as input for SMARTer Stranded Total RNA-Seq Kit v2 – Pico Input Mammalian for fragmentation (with slight modification) and first strand cDNA synthesis followed by addition of illumina adaptors and indexes via 5 cycles of PCR. Indexed libraries were purified using JetSeq SPRI magnetic beads (Bio, # 68031). To remove rRNA fragments the cDNA was cleaved by ZapR with the libraries hybridized to R-probes. The resulting library fragments that were ribo-depleted were further enriched by 16-18 cycles of PCR. The final ribo-depleted libraries were purified using JetSeq SPRI magnetic beads (Bio, # 68031). The concentration of the libraries was measured using Qubit fluorometer (Thermo Fisher Scientific, MA, USA) and then analyzed on Agilent 2100 Bioanalyzer for fragment size distribution.

### RNA seq data analysis

The transcriptome analysis was performed by processing the raw data for removal of low-quality data and adaptor sequences. The raw reads were processed using FastQC for quality assessment and pre-processing which includes removing the adapter sequences and low-quality bases (<q30) using TrimGalore. The high-quality reads were considered for alignment with reference genome using a splice aligner. Only the high-quality data was aligned to reference (Homo sapiensGRCh38.p13 build genome downloaded from Ensembl database) using Hisat2 with the default settings to identify the alignment percentage. Reads were classified into aligned and unaligned reads depending on whether they aligned to the reference genome. Hisat2 was used to calculate raw read counts for each gene after alignment. For differential gene expression analysis, low counts genes with a mean raw read count < 100 per subset were excluded. All differential gene expression analysis was performed using R package DESeq2. Genes with Benjamini-Hochberg (B-H) adjusted p-value ≤ 0.05 were considered significant. Further, genes with Log2 fold change of greater than or equal to +1 were considered upregulated whereas less than or equal to -1 were considered downregulated genes. For heatmaps, log normalized counts transformed to their Z-scores were used. Z-score was calculated by subtracting normalized counts with the mean of the entire dataset and then dividing it by the standard deviation (σ) of the dataset. Ward.D2 method was used for hierarchal clustering which employs the sum of squares of Euclidian distances to perform clustering. R packages clusterProfiler, ggplot2 and dplyr were used for data transformation and plotting.

### Pathway enrichment analysis

KEGG enrichment analysis was performed using the R package clusterProfiler which supports downloading the latest online version of KEGG data from its website. Pathway enrichment query was run using genome wide human annotation (org.Hs.eg.db), B-H adjusted p-value cut-off ≤ 0.05 and minimum set size > 10.

### Statistical analysis

The data was curated in MS Excel and statistical analysis was performed on GraphPad prism and by using R programming language. For analysis of groups, unpaired two-tailed t test was used and p values were interpreted as * p≤0.05;** p≤0.01; *** p≤0.001, ****p<0.0001.

## Results

### Characterization of the CD8 T cell responses in dengue patients

Surface markers HLADR and CD38 are well-defined for identification of activated and expanding CD8 T cells (19, 22, 23). Previous studies including ours in children with confirmed dengue showed that the HLADR^+^CD38^+^ CD8 T cells expand massively during the febrile phase. Therefore, we first asked what was the expansion of the HLADR^+^CD38^+^ CD8 T cell population in dengue confirmed adult acute febrile patients **(Table 1)**. Consistent with previous studies (19), we found that the HLADR^+^CD38^+^ CD8 T cell population expand massively in these patients as compared to healthy **(Figure 1A)** with frequencies reaching as high as 80% of the total CD8 T cells **(Figure 1B, left)** and the absolute numbers reaching as high as a million cells / ml of the blood **(Figure 1C, right)**. Also consistent with previous studies in children (19), these massively expanding HLADR^+^CD38^+^ CD8 T cell population were blasting as seen by high forward scatter (FSC) and side scatter (SSC) and robustly expressed proliferating cell marker (Ki67), downregulated markers of naïve cells (CD45RA, CCR7 and CD127), upregulated inflammatory homing receptors (CX3CR1, CCR5) and upregulated several markers indicative of cytotoxic T-cell effector functions (granzyme A, granzyme B and granulysin). Interestingly, only a small proportion of these massively expanding HLADR^+^CD38^+^ CD8 T cells made IFNγ when stimulated *in vitro* with peptide pool spanning the entire dengue proteome (**Figure 1E)**. The highest frequency of the IFNγ+ cells among the gated HLADR^+^CD38^+^ CD8 T population was 5% with mean ± SEM value being 0.75 ± 0.18. While the total HLADR^+^CD38^+^ CD8 T cell absolute numbers reached up to a maximum of 1×10^6^ cells /ml blood, the total IFNγ producing HLADR^+^CD38^+^ CD8 T cell absolute numbers reached only up to a maximum of 5×10^4^ cells /ml blood with mean ± SEM: being 4.266 ×10^3^ ± 1.580 × 10^3^. Nearly a third of the HLADR^+^CD38^+^ CD8 T cells expressed CD69 *ex vivo* **(Fig 1H, left)**. After peptide stimulation, the IFNγ producing cells that have evolved were found among these CD69^+^ cells but the IFNγ producing population constituted only a small fraction of these CD69^+^ cells **(Figure 1H, middle and right)**. Taken together, these results show that while the HLADR^+^CD38^+^ cells CD8 T cells have a highly differentiated effector phenotype and expand massively with a substantial portion expressing CD69 ex vivo, only a small proportion of them make IFNγ in response to *in vitro* stimulation with dengue peptides.

**Table 1:** Summary of dengue patients analyzed in this study

|  |  |
| --- | --- |
| Total no. of patients | 100 |
| No. of males/females | 57/43 |
| Age (yr) [range (avg)] | 25.5 (15-66) |
| No. of days post-onset of clinical symptoms<br>[range (avg)] | 3-12 (7.5) |

**Figure 1.**
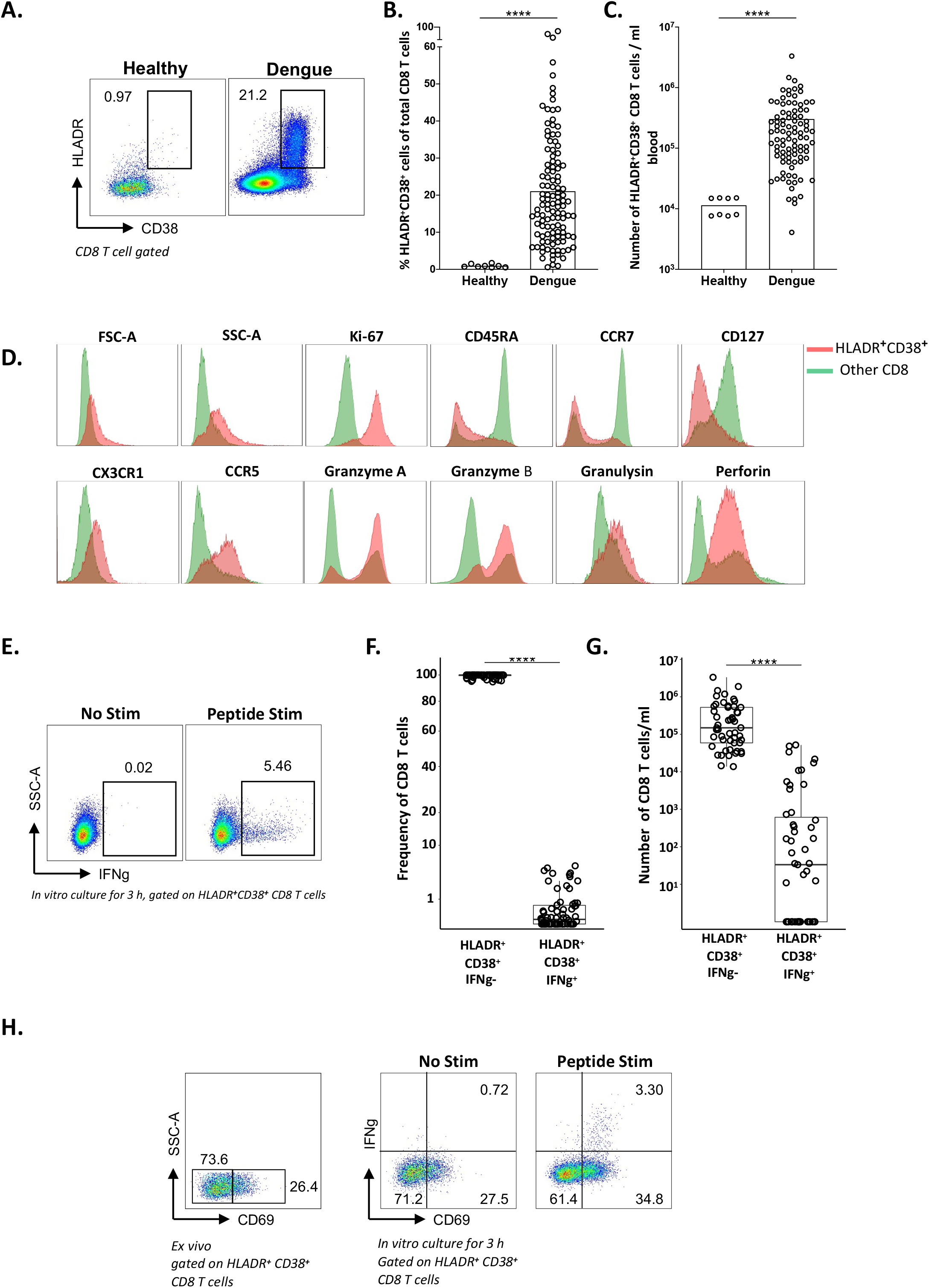
Analysis of the CD8 T cell responses in acute dengue febrile patients: **(A)** Example of flow cytometry plot gated on total CD8 T cell showing evaluation of the HLADR^+^CD38^+^ CD8 T cells in healthy (left) and dengue febrile patient (right). **(B)** The scatter plots show the frequencies (percentages) of HLADR^+^CD38^+^CD8 T cells among the gated CD8 T cell population and **(C)** absolute numbers per millilitre of blood in healthy and dengue febrile adults. **(D)** The histogram plots represents the frequency of various phenotypic markers studied. HLADR^+^CD38^+^ CD8 T population is red color and all other CD8 T cells green. **(E)** Example of flow cytometry plots showing IFNγ producing HLADR^+^CD38^+^CD8 T cells in unstimulated and peptides stimulated dengue patient. **(F)** The scatter plots showing frequencies and **(G)** numbers of non-IFNγ producing and IFNγ producing HLADR^+^CD38^+^CD8 T cells after peptide stimulation in acute dengue patients. **(H)** Flow cytometry plot shows ex-vivo expression of CD69 on HLADR^+^CD38^+^ CD8 T cells (right) and in-vitro expression of CD69 and IFNγ on HLADR^+^CD38^+^ CD8 T cells with and without dengue peptide stimulation (left). In B, C, F and G each dot represents analysis of different individual. Bar represents the mean. Significance values were calculated using a two-tailed Mann Whitney’s U test and are indicated by: ****, p≤0.0001.

### RNA sequencing of functionally distinct HLADR^+^CD38^+^ CD8 T cells

To address what other cytokines/ chemokines do these massively expanding HLADR^+^ CD38^+^ CD8 T cells express upon peptide stimulation and what transcriptional profiles distinguish the CD69^+^IFNγ^+^, CD69^+^IFN-γ^-^, and CD69^-^IFN-γ^-^ subsets, we stimulated PBMC of three individual patients **(Supplementary Table 1)** with dengue peptides for 3 h *in vitro* and then sorted the CD69^+^IFNγ^+^, CD69^+^IFNγ^-^, and CD69^-^IFNγ^-^ subsets among the gated HLADR^+^CD38^+^ CD8 T cell population followed by RNA seq analysis. Because the RNA analysis cannot be performed after intracellular staining, we employed IFNγ capture assay for identifying and sorting the CD69^+^IFNγ ^+^, CD69^+^IFNγ^-^, and CD69^-^IFNγ^-^ subsets. As a control, we also sorted two other subpopulations from samples that were *in vitro* cultured in the absence of peptide stimulation from four patients. These include the total HLADR^+^CD38^+^ CD8 T cell (here after referred to as unstimulated double positive or unstim DP) and the HLADR^-^CD38^-^ CD8 T cell populations (hereafter referred to as unstimulated double negative or DN). **Figure 2A**, shows the gating strategy. The post sort purity was >96%.

**Figure 2.**
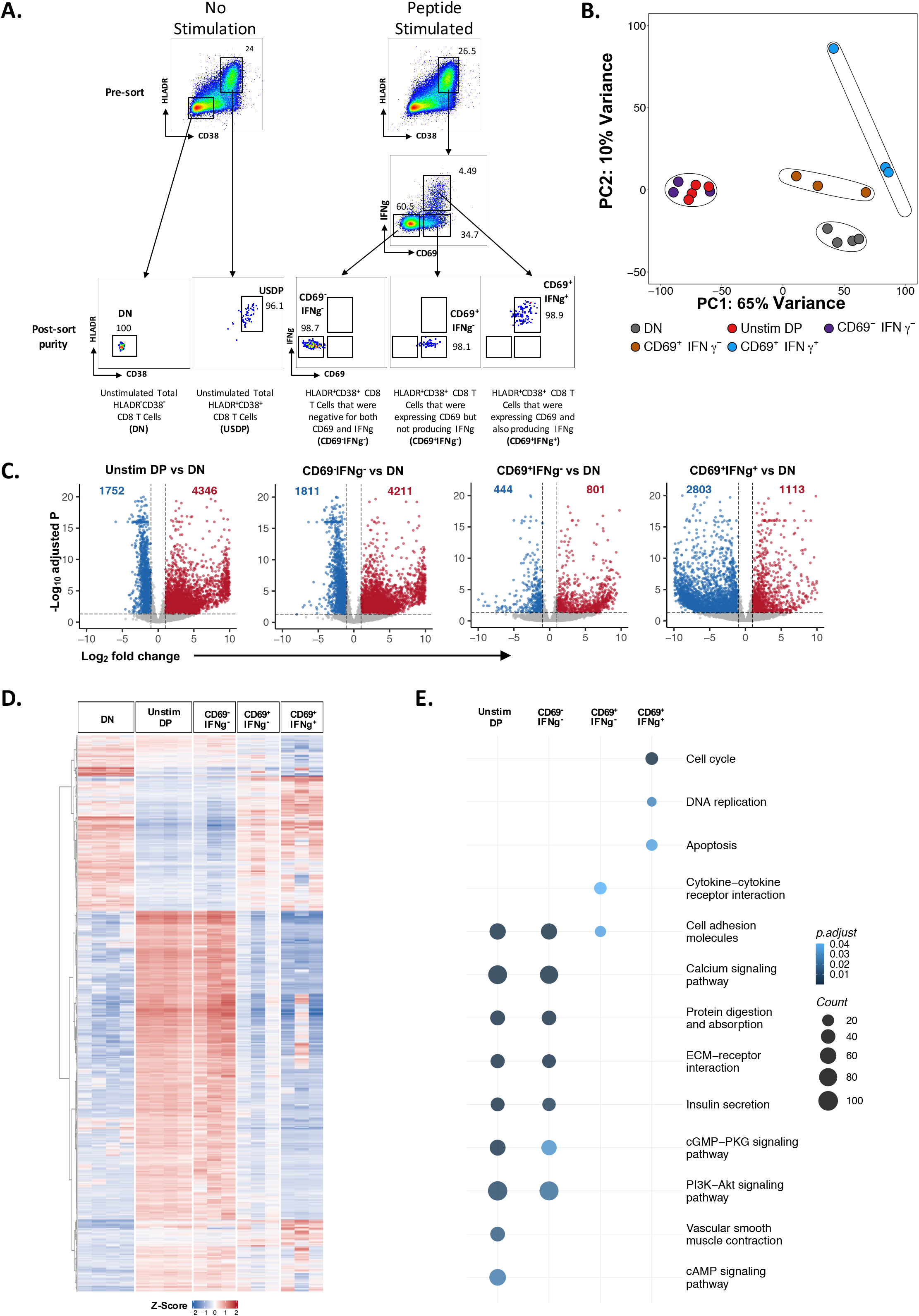
Sorting of IFNγ producing HLADR^+^ CD38^+^ CD8 T CD8 T cells and other activated CD8 T cells from dengue patients and their global transcriptional profiling / analysis using RNA-seq. **(A)** The subsets of activated CD8 T cells were sorted using flow-cytometry based on the expression of CD38, HLADR, CD69 and IFNγ following ex-vivo stimulation for 3 hours with and without dengue peptides. The sorting strategy and purity of sort is shown. **(B)** Principal component analysis (PCA) of five subsets, unstimulated HLADR^-^ CD38^-^ (DN, n= 4, grey), unstimulated HLADR^+^ CD38^+^ (Unstim DP, n= 4, red), stimulated HLADR^+^CD38^+^ CD69^-^IFNγ^-^ (CD69^-^ IFNγ^-^, n = 3, violet), stimulated HLADR^+^CD38^+^ CD69^+^IFNγ^-^ (CD69^+^IFNγ^-^, n = 3, brown) and stimulated HLADR^+^CD38^+^ CD69^+^IFNγ^-^ (CD69^+^IFNγ^+^, n = 3, blue) based on 14,959 genes. **(C)** Volcano plots highlighting differentially expressed genes in following comparisons: Unstim DP v/s DN (first from left), CD69^-^IFNγ^-^ v/s DN (second from left), CD69^+^IFNγ^-^ v/s DN (third from left) and CD69^+^IFNγ^+^ v/s DN (right). Each dot represents a gene with log2 fold change (Log^2^FC) on x-axis and negative log10 Benjamini-Hochberg (B-H) adjusted p-value (-Log^10^ adjusted P) on y-axis. Upregulated genes (Log^2^FC > 1 and adjusted p-value < 0.05) are shown in red, downregulated genes (Log^2^FC < -1 and adjusted p-value < 0.05) are shown in blue and genes which are not significant are shown in grey. **(D)** Heatmap showing hierarchal clustering of all DEGs from the comparisons shown in **(C)** based on Ward.D2 algorithm. Normalized gene expression was converted into Z-scores for plotting. **(D)** KEGG pathway enrichment of genes overexpressed in Unstim DP, CD69^-^IFNγ^-^, CD69^+^IFNγ^-^ and CD69^+^IFNγ^+^ with respect to DN is shown. Only Selected pathways amongst significant ones (B-H adjusted p-value < 0.05) are shown. For each pathway, the intensity of the colored dot represents B-H adjusted p-value and size represents the number of overexpressed genes mapped.

The DEG’s found in the unstim DP in this study was largely consistent with the DEGs with our previous microarray analysis wherein we compared *ex vivo* sorted total HLADR^+^CD38^+^ CD8 T cells from dengue febrile children versus CD45RA^+^CCR7^+^ naïve CD8 T cells from the same patients (19) (**Supplementary Table 2)**. This result suggested that the transcriptional profiles of the unstim DP that were obtained after *in vitro* culture are similar to the transcriptional profiles of the *ex vivo* sorted DP.

Principal component analysis based on 14,959 differentially expressed genes (DEGs) in one or more subsets compared to the DN subset **(Supplementary Table 3)** showed that while the DN subset formed a separate cluster from all the other subsets, the unstim DP and the stimulated CD69^-^IFNγ^-^ subsets formed an overlapping cluster whereas the stimulated CD69^+^IFNγ^+^ and CD69^+^IFNγ^-^ subsets formed distinct clusters (**Figure 2B)**. This result indicated that while there were no major gene expression differences between the unstim-DP and the stimulated CD69^-^ IFNγ^-^ subsets, there were major gene expression differences between the unstim-DP / stimulated CD69^+^IFNγ^+^ subsets, the stimulated CD69^+^IFNγ^-^subset, and the stimulated CD69^-^IFNγ^-^ subset. Consistent with this, the DEGs found in the Unstim DP and the stimulated CD69^-^IFNγ^-^ subsets were similar; whereas the DEGs found in the stimulated CD69^+^IFNγ^-^ subset and the stimulated CD69^+^IFNγ^+^ subsets were distinct in numbers **(Fig 2C)**, expression levels **(Fig 2D and Suppl Table 3A)** and the significant biological pathways associated **(Fig 2E)**. While some biological processes such as cell cycle / DNA replication and apoptosis were significantly upregulated in the CD69^+^IFNγ^+^ subset, processes such as cytokine-cytokine receptor interaction were preferential to the CD69^+^IFNγ^-^ subset compared to the DN (**Fig 2E**). By contrast, both the unstim DP as well as the stimulated CD69^+^IFNγ^-^ subsets showed preference to some other processes related to calcium signaling, protein digestion/ absorption, extra cellular matrix interaction, insulin secretion, cGMP-PKG and PI3K-AKT signaling. On the other hand, the unstim-DP showed preference to processes related to vascular smooth muscle contraction and cAMP signaling. Because this pathway analysis did not account for qualitative and quantitative differences between the subsets, we performed a side-by-side comparison of relative expression of select genes of interest related to cytokines/ chemokines and their receptors, cytotoxic effector functions, TCR signaling, co-stimulation, proliferation, protein translation, metabolism, transcriptional machinery and other T cell effector functions. The notable findings are elaborated in the sections below.

### Differences in the expression of cytokines/chemokines and their receptors in the three HLADR^+^CD38^+^ CD8 T cell subsets

The relative expression of the DEGs related to cytokine/ chemokines is shown in **Fig 3A and supplementary Table 4**. Notably, we found that the CD69^+^IFNγ^+^ subset robustly expressed two other chemokines (XCL1 and XCL2) in addition to the anti-viral cytokine IFNγ. There was very little or no expression of any of these cytokines/ chemokines in the DN, unstim-DP, CD69^-^IFNγ^-^ and the CD69^+^IFNγ^-^ subsets. On the other hand, both the unstim-DP and the CD69^-^IFNγ^-^ subset showed a bias towards expression of CXCL13, IL-16, IL-17B, IL-19, IL-34, IL-37, IL-36RN, PDGFB, TGFA and GRN.

**Figure 3.**
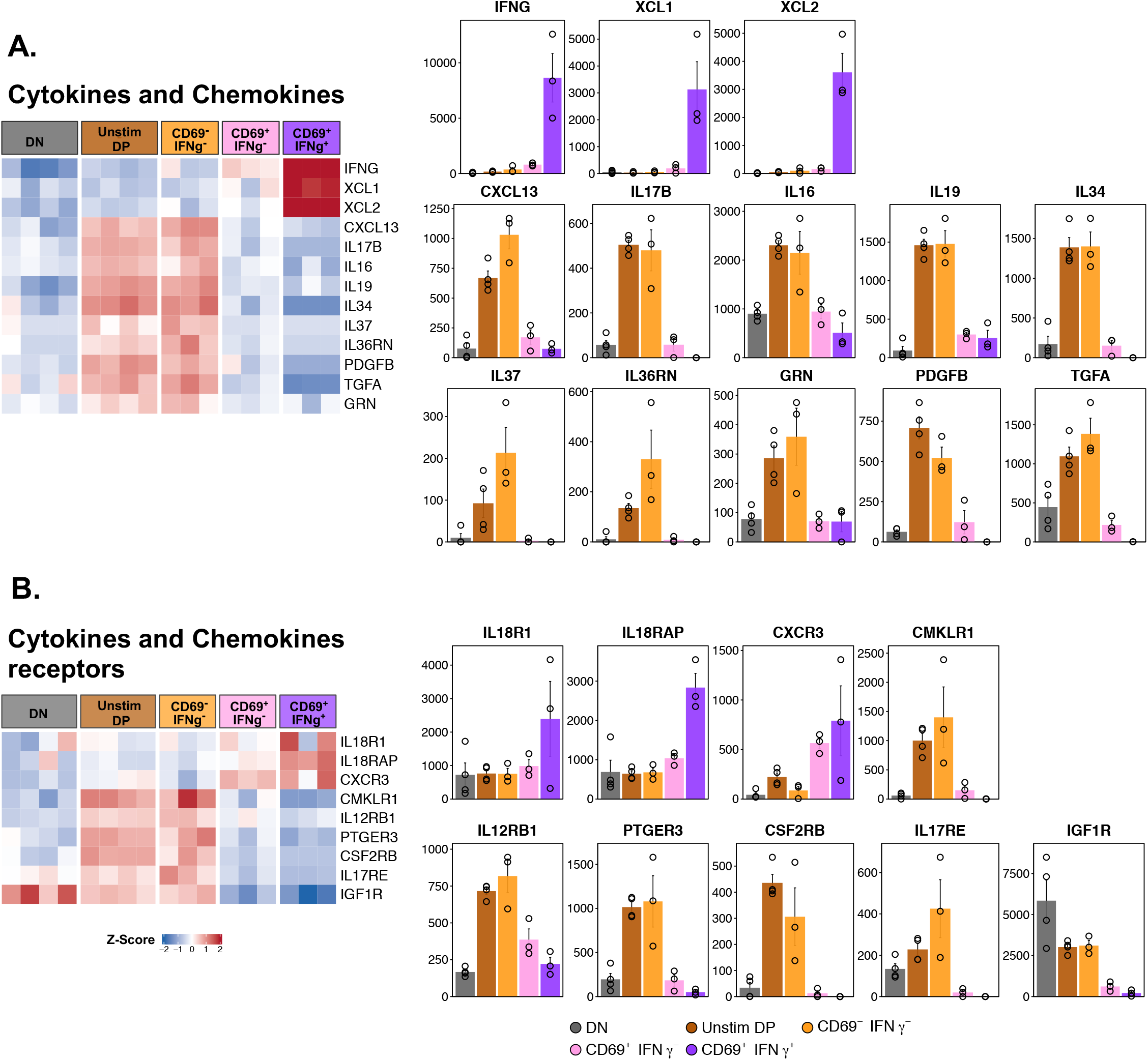
Key genes of interest differentially regulated in CD69^-^IFNγ^-^ subset when compared with CD69^+^IFNγ^-^ and CD69^+^IFNγ^+^ subsets. Heatmap showing the expression profile of key genes of interest (left). Color gradient represents the normalized gene expression transformed into z-scores from blue (low expression) to red (high expression). For each gene, bar plots representing average normalized counts of DN, Unstim DP, CD69^-^IFNγ^-^, CD69^+^IFNγ^-^ and CD69^+^IFNγ^+^ subsets are shown. Error bar represents standard error (SEM).

Taken together, these results show that the CD69^+^IFNγ^+^ subset robustly expressed key genes involved in anti-viral functions/ MHC class-I upregulation (IFNγ), dendritic cell cross talk (XCL1) (24) and migration of cells during the immunological responses (XCL2) (25). By contrast, the CD69^-^IFNγ^-^ subset expressed several other cytokines/chemokines that had other immunological relevant functions. Noteworthy amongst these were - Chemokine CXCL13 that is associated with CD8 T cell dysfunction (26); cytokines such as IL-16, that has been shown to inhibit IL-2 production and is associated with CD4 dysfunction in HIV (27); IL-17b, that promotes survival and proliferation (28); IL-19 that downregulates CTL responses in helminth infections (29); IL-34, that mediates transplant tolerance (30); IL-37, that is a potent anti-inflammatory cytokine and inhibits trained immunity (31); IL-36RN, that inhibits activation of NFkB (32); PDGFb, that promotes proliferation and survival but depresses production of IL-4, IL-5 and IFNγ (33); TGFα, that acts in synergy with TGFα to also promote proliferation and survival (34). We also observed high levels of GRN, which is a Granulin that is a key lysosomal protein that contributes to inflammation and enhances that formation of inducible Tregs (iTregs) that secrete IL-10 (35) **(Figure 3A, Supplementary Table 4)**.

We also observed an increased expression of IL-18R1, IL-18RAP and CXCR3 in both the CD69^+^IFNγ^-^ and the CD69^+^IFNγ^+^ subsets. IL-18R1 is a receptor for the cytokine IL-18 that mediates IFNγ synthesis (36); IL-18RAP, is an accessory subunit of the IL-18 receptor that leads to the activation of NFkB during cell mediated immunity (37) and CXCR3, a receptor for C-X-C chemokines CXCL9, CXCL10 and CXCL11 that is known to be upregulated on activated effector CD8 T cells and mediates survival and proliferation (38). By contrast, the CD69^-^IFNγ^-^ subset showed better expression of cytokine receptors such as CMKLR1, a receptor for chemoattractant chemerin that augments T cell mediated cytotoxicity (39); IL-12RB1, that is a receptor for pro-inflammatory cytokines IL-12 and IL-23 (40); PTGER3, a receptor for prostaglandin E2 that is required for development of fever and has been shown to be critical for IL-17 driven inflammation (41); CSF2RB/IL-15RB1 that is a common beta chain for cytokines IL-3, IL-5 and is regarded as a marker for antigen stimulated cells (42); IL-17RE, that is a receptor for pro-inflammatory cytokine IL-17C (28) and IGF1R, a high affinity receptor for IGF1 and results in the activation of Pi3K-AKT and Ras-MAPK pathways (43) **(Figure 3B, Supplementary Table 5)**.

Taken together, these data show that the CD69^+^IFNγ^+^, CD69^+^IFNγ^-^ and CD69^-^IFNγ^-^ subsets differ in expression of several cytokines, chemokines and their receptor genes that have immunological relevance and distinct functions.

### Differences in the expression of genes related to cytotoxic effector functions in the HLADR^+^CD38^+^ CD8 T cell subsets

The most striking feature of CD8 T cells is their ability to kill virally infected CD8 T cells through their cytotoxic functions. Therefore, we next asked which of the subsets of HLADR^+^CD38^+^ CD8 T cells did the cytotoxic gene signature filter. We observed that several key genes associated with cytotoxic effector functions were upregulated in both CD69^+^IFNγ^-^ and CD69^+^IFNγ^+^ subsets, with the latter having more robust expression. Notable were the well-known genes involved in cytotoxic effector functions such as granzyme B (GzmB), granzyme A (GzmA), granzyme H (GzmH), perforin (PerF1), Granulysin (Gnly), Cathepsin W (CTSW); CRTAM, a gene known to promote NK cell cytotoxicity and CTL function in CD8 T cells (44); and IGF2R, a gene responsible for intracellular trafficking of lysosomal enzymes (45). By contrast no genes related to cytotoxicity filtered in the CD69^-^IFNγ^-^ subset **(Figure 4A, Supplementary Table 6)**. Taken together, these results indicate that both the CD69^+^IFNγ^+^ subset as well as the CD69^+^IFNγ^-^ subset are highly enriched for expression of key genes involved in cytotoxic effector functions.

**Figure 4.**
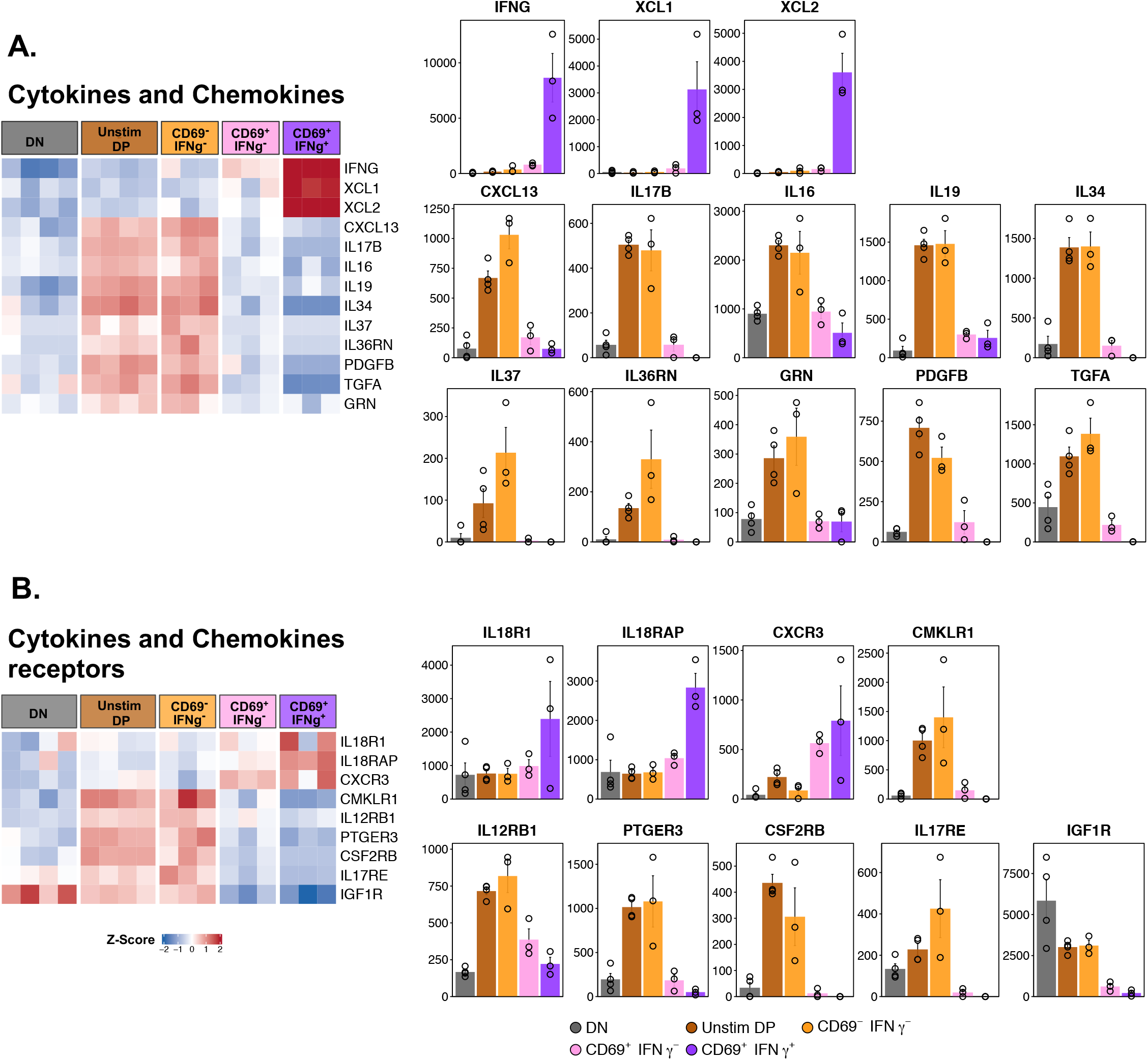
Notable genes upregulated in CD69^+^IFNγ^-^ and CD69^+^IFNγ^+^ subset. Heatmap showing the expression profile of notable genes which were upregulated in CD69^+^IFNγ- and CD69^+^IFNγ^+^ subset as compared to DN (left). Genes are segregated based on their functions. Color gradient represents the normalized gene expression transformed into z-scores from blue (low expression) to red (high expression). For each gene, bar plots representing average normalized counts of DN, Unstim DP, CD69^-^IFNγ^-^, CD69^+^IFNγ^-^ and CD69^+^IFNγ^+^ subsets are shown. Error bar represents standard error (SEM).

### Differences in the expression of genes involved in TCR signaling / co-stimulation in the HLADR^+^CD38^+^ CD8 T cell subsets

Consistent with the cytotoxic and or cytokine effector functions, the CD69^+^IFNγ^+^ and the CD69^+^IFNγ^-^ subsets also showed robust expression of several key genes involved in TCR signaling and co-stimulation. Notable among these include TNFSF9/4-1-BBL, that is a ligand for TNFRSF9/4-1-BB (46); ICOS/CD278, that enhance T cell responses (47); HAVCR1/TIM-1, a P-selectin that regulates the expression of a number of cytokines including IL-10 (48). While JAK2 and SH2D2A that participate in T cell signaling were upregulated, balancing these, both these subsets also showed expression of inhibitor receptors Lag 3 and CTLA-4 (49); PTPN22 (50) and HAVCR2 (51), that downregulate TCR signaling; and, phosphatases such as DUSP4, that downregulate MAPK and interleukin signaling (52). Additionally, the CD69^-^IFNγ^+^ also preferentially expressed the co-stimulatory molecule TNFSF14 (53).

By contrast, the CD69^-^IFNγ^-^ subset was biased towards expression genes encoding a different set of costimulatory molecules that regulate T cell responses, CD276 (54) and CD80 (55); genes belonging to the tyrosine phosphatase family – PTPRM, that is required for cell growth (56); and PTPRF that is known to negatively regulate TCR signaling by dephosphorylation of LCK and Fyn (57). Interestingly, we also observed upregulation of genes that are required for processes important to dengue disease - Endothelin 1 (EDNRA) that plays a role in vasoconstriction (58), Claudin 5 and 6 (CLDN5 and CLDN6) that are components of tight junctions and are required for extravasation of immune cells (59) **(Figure 4B, Supplementary table 7)**.

### Differences in the expression of genes involved in transcription, translation, metabolism, and proliferation in the CD69^+^IFNγ^+^ subset

A careful comparison of the CD69^+^IFNγ^+^ subset against all the four other populations sorted revealed 70 genes that were specifically and uniquely upregulated only in the cells that were capable of making IFNγ in response to dengue peptides **(Figure 5, and Supplementary Table 8)**. Of these were genes associated with proliferation, TCR signaling, inflammation, energy metabolism and protein translation **(Figure 5A)**. Notably, genes that participate in DNA replication and proliferation; CTPS1, an enzyme responsible for conversion of UTP to CTP (60); ODC1, a rate limiting enzyme in the polyamine biosynthesis pathway (61); PAICS, a key enzyme in purine biosynthesis and PPAT, also another enzyme that is of the phosphoribosyltransferase family (62) and TFRC / CD71, a transferrin receptor known to be highly upregulated on activated and proliferating T and B cells(19, 63) **(Figure 5B, Supplementary Table 8)**. It is interesting to note a recent study showed that expression of CD71 may also mark memory precursor CD8 T cells (23).

**Figure 5.**
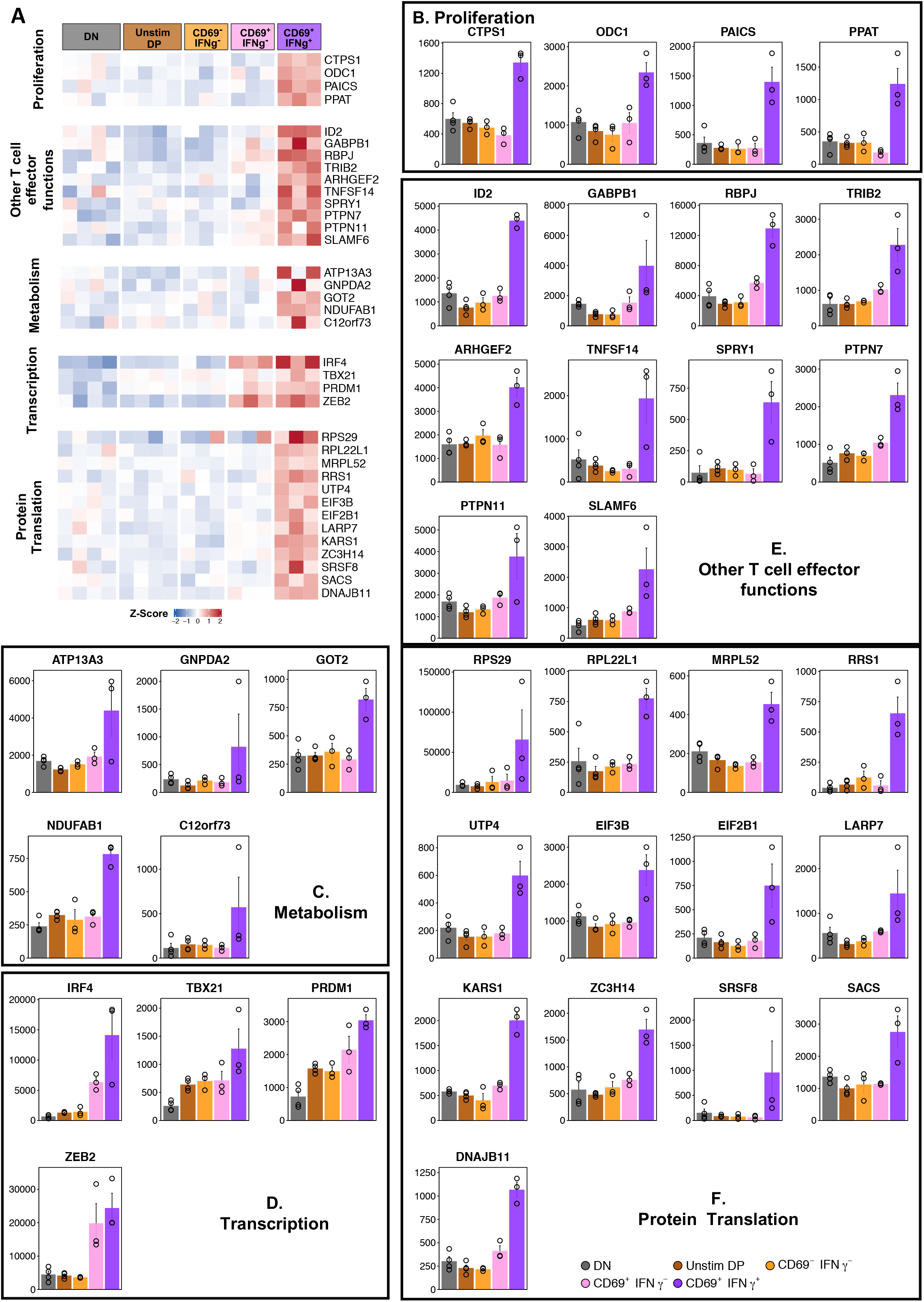
Genes uniquely overexpressed in CD69^+^IFNγ^+^ subset categorized by function. Heatmap showing the expression profile of genes uniquely overexpressed in CD69^+^IFNγ^+^ subset (left). The genes are categorized into putative functions such as DNA replication and proliferation, metabolism, T cell functions and protein translation. Color gradient represents the normalized gene expression transformed into z-scores from blue (low expression) to red (high expression). For each gene, bar plots representing average normalized counts of DN, Unstim DP, CD69^-^IFNγ^-^, CD69^+^IFNγ^-^ and CD69^+^IFNγ^+^ subsets are shown. Error bar represents standard error (SEM).

These CD69^+^IFNγ^+^ HLADR^+^CD38^+^ CD8 T cells were also highly metabolically active as indicated by the distinct expression of genes such as APT13A3, a p-type ATPase; GNPDA2, an enzyme that converts glucosamine-6-phosphate to fructose-6-phosphate (64); GOT2, is known to be expressed in activated CD8 T cells and is required for amino acid metabolism and TCA cycle (65); NDUFAB1 and C12orf73, both involved in the mitochondrial respiratory chain complex (66) **(Figure 5C, Supplementary Table 8)**. As enhanced cellular metabolism is an indicator for cell survival, it again suggests that the cells that are capable of making IFNγ in response to dengue peptides are probably the lineage that survives and forms the memory CD8 T cell pool.

We also observed key genes of interest that are necessary for T cell function, such as, transcription factors ID2, that promotes survival and differentiation of CD8 T cells (67); GABPB1, an Ets transcription factor, shown to be critically responsible for antigen-stimulated T cell responses (68). Other genes associated with T cell function such as RBPJ, that augments notch signaling (69); TRIB2, involved in MAPK signaling (70); ARHGEF2, shown to promote IL-6 and TNFα secretion (71); TNFSF14/LTg (53), that promotes co-stimulation and the SPRY1 (72), that has a dual effect on cytokine production depending on the state of the T cell were found to be robustly expressed. Interestingly, we also observed that the CD69^+^IFNγ^+^ HLADR^+^CD38^+^ CD8 T cells also expressed genes that probably regulate overabundance of TCR mediated immune activation. Phosphatases such as PTPN7 and PTPN11 that downregulate TCR signaling were increased (57), along with SLAMF6, that is a known immune checkpoint inhibitor (73) **(Figure 5D, Suppl Table 8)**.

Lastly, lineage commitment transcription factors of IRF4, Tbx21 (T-Bet), Prdm1 (Blimp1) and Zeb2 that are well known to drive cytotoxic T cell responses (74, 75) while being upregulated in both CD69^+^IFNγ^+^ and CD69^+^IFNγ^+^, the IFNγ producers had significantly higher expression as compared to the CD69^+^IFNγ^-^ subset indicating that this subset was more driven towards classical cytotoxic T cell responses **(Figure 5E, Supplementary Table 8)**.

A striking observation we made in the CD69^+^IFNγ^+^ HLADR^+^CD38^+^ CD8 T cells was the expression of a number of genes that were involved in protein translation such as components of the ribosome RPS29, RPL22L1 and MRPL52; ribosome biogenesis regulators such as RRS1 and UTP4; translational initiation factors EIF3B, EIF2B1 and LARP7; KARS1, a lysyl-tRNA-synthetase; ZC3H14, critically required for mRNA stability; SRSF8, required for mRNA splicing and protein chaperones SACS and DNAJB11 were all found to be highly expressed **(Figure 5F, Supplementary Table 8)**.

Taken together, this data shows that the CD69^+^IFNγ ^-^ subsets were also enriched for expression of several key genes that contribute to proliferation, T cell effector function, expression of lineage commitment transcription factors and processes such as protein translation that perhaps facilitates the higher expression of genes involved in immune responses, including IFNγ.

### Differences in the expansion of the HLADR^+^CD38^+^ CD8 T cell subsets depending on disease severity

We next wondered whether the expansion of the total HLADR^+^ CD38^+^ CD8 T cell population or the IFNγ producing subset among these differ between the SD, DW and DI cases **(Suppl Table 9)**. Enumeration of frequency and numbers of HLADR^+^CD38^+^ CD8 T cells **(Figure 6A)**, or IFNγ^+^ HLADR^+^CD38^+^ CD8 T cells **(Figure 6B)** showed no statistically significant differences between the SD, DW and DI cases.

**Figure 6.**
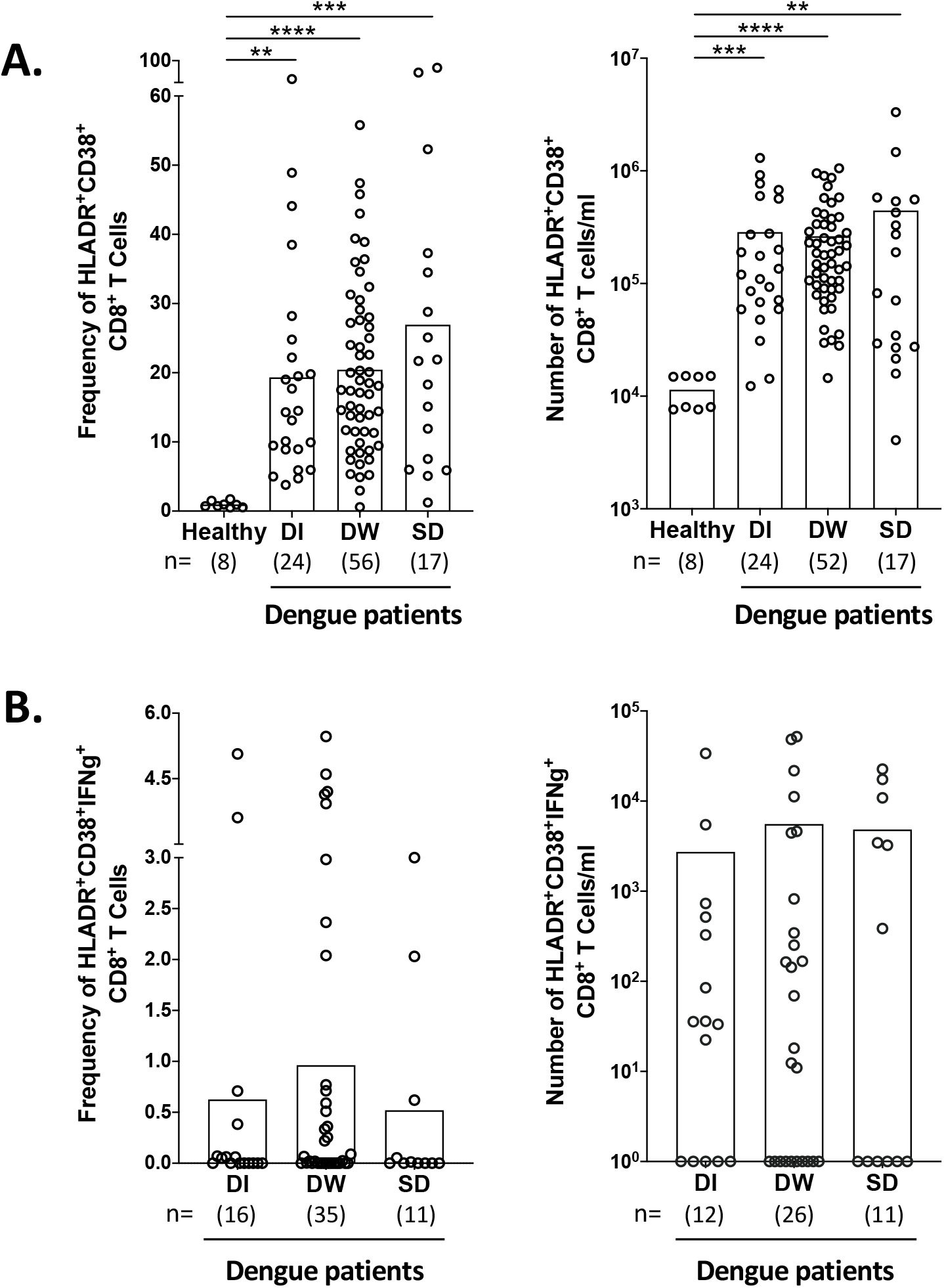
Comparison of CD8 T Cell response in dengue patients depending on disease severity. **(A)** The scatter plots shows frequencies (left) and number (right) of HLADR^+^CD38^+^ cells among the gated CD8 T cell population in acute dengue patients classified according to different grades of disease severity. **(B)** The scatter plots showing frequencies (left) and number (right) of IFNγ producing HLADR^**+**^CD38^**+**^CD8 T cells after peptide stimulation in acute dengue patients classified according to different grades of disease severity. Bar represents the mean. Significance values are calculated using Kruskal Wallis one-way analysis of variance (ANOVA) followed by Dunn’s multiple comparison test. The notation ns means not significant, with p > 0.05.

## Discussion

In summary, we show that a discrete population of the HLADR and CD38 expressing CD8 T cells with a highly differentiated effector phenotype expand massively during the acute febrile phase of dengue natural infection. This is consistent with previous studies. Although about a third of these massively expanding HLADR^+^ CD38^+^ CD8 T cells found in the peripheral blood were of CD69^high^ phenotype, only a small fraction of them produced IFNγ upon *in vitro* stimulation with peptide pools spanning the entire dengue proteome. We found that while these small proportion of CD69^+^IFNγ^+^ subset expressed key genes involved in protein translation, cellular metabolism, proliferation and dendritic cell cross talk; both the CD69^+^IFNγ^+^ and larger proportion of CD69^+^ IFNγ^-^ subsets expressed key genes that are aligned to an antigen responsive phenotype and are involved in cytotoxic effector functions, regulation of T cell receptor signaling, regulation of signaling by MAP kinases/ IL-18 and other growth factors, chemotaxis and T cell trafficking to inflamed tissues with the expression being highest in the CD69^+^IFNγ^+^ subset. Additionally, we show that the CD69^-^IFNγ^-^ subset which is the bulk of the HLADR^+^CD38^+^ population expressed key cytokine/ chemokine genes that are typically implicated in dampening the immune response including those that are associated with CD8 T cell dysfunction, inhibition of IL-2 production, IL-25 signaling, NFKB activation, IL-4, IL-5, IFNγ production; down regulation of T cell responses, and promoting the formation of the inducible T-regs. The expression of these immunoregulators in the CD69^-^IFNγ^-^ subset is likely to be constitutive and unlikely to require *in vitro* peptide stimulation since the expression of these same genes was also observed in the unstimulated HLADR^+^CD38^+^ CD8 T cells (of which, nearly third of the cells are likely to be CD69^-^IFNγ ^-^ phenotype). In the peripheral blood, neither the expansion of the total HLADR^+^ CD38^+^ CD8 T cells nor the expansion of the IFNγ producing population among these was significantly different between the SD, DW and DI cases. These findings provide important baseline information for future studies to interrogate the tissue location and dynamics of these different cell subsets during the transition from acute febrile phase to memory / recall responses in dengue natural infection and or vaccination.

Previous studies in human hepatitis virus or cytomegalovirus natural infections and influenza/ yellow fever live attenuated vaccination showed that a substantial fraction of the antigen specific CD8 T cells that expand in response to viral infection are of stunned phenotype as assessed by their inability to produce IFNγ upon *in vitro* peptide stimulation (19, 76-78). A recent study showed that those antigen-specific CD8 T cells that are capable of expressing the IFNγ gene upon *in vitro* peptide stimulation reliably co-express chemokines such as XCL1 and XCl2 (44). Our result showing co-expression of XCL1 and XCl2 among the IFNγ producing cells is consistent with these previous studies and suggests that, perhaps, the cytokine functional response properties of the CD8 T cells in dengue natural infection is likely to be similar to these the other human viral infections.

Interestingly, we found that the CD69^+^IFNγ^+^ subset, in addition to expressing the IFNγ, XCL1, XCL2 and other genes associated with cytotoxic effector functions, also upregulated several other genes that contribute to an enhanced proliferation, protein translation metabolic cellular programs involving mitochondrial respiratory chain. In this perspective, it is important to note that the development of memory T cell development involves expression of large number of metabolism related genes (23). Our results, together with these previous studies, raises an interesting question on whether the memory precursors are present in only the CD69^+^IFNγ^+^ subset which constitutes a small fraction of the massively expanding HLADR^+^CD38^+^ CD8 T cells in dengue infection whose frequencies are similar to what is expected to survive after infection is resolved (79-81). This is more likely to be the case given the observations from a recent study in people vaccinated with live attenuated dengue vaccine showing that the TCR clonotypes found among the IFNγ producing CD8 T cells was similar at the peak of the CD8 expansion and memory phase (23). Similar studies are needed in unvaccinated and vaccinated people following dengue natural infection to understand the precursors of memory development in dengue natural infection.

Our data shows that the expansion of the HLADR^+^CD38^+^ CD8 T cells or the IFNγ producing cells did not significantly differ between the SD versus, DW versus DI cases. This finding is consistent with a recent study showing that the expansion of the IFNγ producing cells was similar between cases with dengue hemorrhage fever (DHF) and dengue infection (DI) (16). Additionally, this recent study also compared transcriptional profiles of the IFNγ expressing cells in patients with different grades of disease severity and found no striking difference. Thus, this study the emerging idea that the CD8 T cells responses may be protective rather than pathological in dengue.

The transcriptional profiles of the IFNγ producing cells that we described in this study are also consistent with the observations from a recent dengue patient study that performed single cell transcriptomics of the IFNγ producing versus the other CD8 T cells after peptide stimulation (16). In addition to confirming these previous studies, here we provide a detailed understanding of the unique and common transcriptional signatures associated with each of the functionally distinct subsets within the massively expanding HLADR^+^CD38^+^ population in the dengue patients (i.e., CD69^-^IFNγ^-^ versus the CD69^+^IFNγ^-^ versus the CD69^+^IFNγ^+^ subsets). These results suggest that there could be distinct functional lineages of CD8 T cells within these highly activated massively expanding HLADR^+^CD38^+^ CD8 T cells. Of these three subsets, the CD69^+^IFNγ^+^ subset can be undoubtedly interpreted as antigen specific since they are responding to *in vitro* peptide stimulation. However, this CD69^+^IFNγ^+^ subset represents only a minor fraction of these massively expanding HLADR^+^CD38^+^ CD8 T cells. Whether the CD69^+^IFNγ^--^subset and the CD69^-^IFNγ^--^subsets (which together form >90% of these massively expanding CD8 T cells) truly represent antigen specific populations or bystander responses remains to be addressed. Interestingly, a recent study by Waickman et al (23), showed that people that were vaccinated with live attenuated dengue virus (TAK-003) also elicit a massive expansion of the HLADR^+^CD38^+^ CD8 T cells but only a small proportion of them make IFNγ upon in vitro stimulation. Thus, studies are warranted to determine whether the massively expanding HLADR^+^CD38^+^ CD8 T cells in the vaccinated individuals are also comprised of the CD69^-^IFNγ^--^, CD69^+^IFNγ^--^, and CD69^+^IFNγ^+-^ subsets as we have discovered in dengue natural infection and whether each of these subsets from the vaccinated individuals express similar transcriptional profiles as what we found in dengue natural infection; and whether the protective efficacy correlate with one or more of these individual subsets of the CD8 T cells remains to be determined.

It is interesting to note that both the CD69^+^IFNγ^--^, and CD69^+^IFNγ^+-^ subsets among these massively expanding HLADR^+^CD38^+^ CD8 T cells shared a number of DEGs that are directly responsible for cytotoxic functions, chemokines and their receptors and co-stimulatory and inhibitory molecules that perfectly align with an antigen-responsive phenotype. These phenotypes were very similar to those observed in single cell analysis of Flu and CMV specific CD8 T cells (44, 82). Additionally, we found that the CD69^+^IFNγ^+^ subset expressed several unique genes. These observations raise the question on whether CD69^+^IFNγ^-^ subset also has any unique gene signature. Analysis of the DEGs that specially filter to this subset revealed e only four genes– SLFN12L, TENT5C, GGA2 and F2R (PAR1) **(Supplementary table 3)**. Of these, PAR1 is a high affinity receptor for activated Thrombin and studies in murine models have shown that PAR1 expression accelerates calcium mobilization, and is involved in cytotoxic T cell function. Further, T cells from PAR^-/-^ mice have dampened cytokine producing ability suggesting that PAR1/F2R might have a role in T cell function (83). However, our data does not provide any clues towards functionality that that could be uniquely attributed to the CD69^+^IFNγ^-^ subset suggesting that there might be some degree of redundancy in the function of PAR1.

Taken together, our study while improving our understanding of CD8 T cell responses in dengue, also sheds light on the diversity of the expanding CD8 T cell population and warrants further studies to understand and examine the precise role of different functional lineages of CD8 T cells in dengue natural infection and or vaccinees.

### Data sharing

The dengue RNA seq dataset is deposited in Gene expression omnibus (GEO) with the accession code: GSE212034. The private token number for RNA seq analysis uploaded to GEO (GSE212034) for this study is ‘idynkucwbvsfjs’

## Acknowledgements

This work was funded by Dept of Biotechnology, Govt of India, grant number: BT/PR28416/MED/29/1310/2018 and U.S. National Institutes of Health grant ICIDR 1UO1A/115654. We thank Aditya Rathi (ICGEB-TACF) for FACS sorting of the PBMCs. We thank Dr. Daniela Weiskopf, La Jolla Institute of Immunology for CD8 megapools. The authors thank Mr. Satendra Singh, Mr. Ajay Singh, ICGEB, New Delhi for technical support.

## Contributors

Experimental work, data acquisition and analysis was performed by P.S., P.B.,E.S.R.,D.M.,K.S.,Y.C.,K.N,S.J.,P.S. Conceptualization and implementation by P.S., P.B.,E.S.R.,D.M.,K.S.,Y.C.,K.N., S.J.,M.S.,N.W.,M.K.K.,A.C. Manuscript writing by P.S., P.B., M.K.K., A.C. All authors contributed to reviewing and editing the manuscript.

## Legends

**Table 1:** Characteristics of dengue confirmed adult acute febrile patients

**Supplementary Table 1:** Characteristics of dengue confirmed adult acute febrile patients from whom DN, Unstim DP, CD69^-^IFNγ^-^, CD69^+^IFNγ^-^ and CD69^+^IFNγ^+^ subsets were sorted.

**Supplementary Table 2:** Comparison of previous microarray analysis of gene signature of HLADR^+^CD38^+^ CD8 T cells and CD45RA^+^CCR7^+^ CD8 T cells from dengue confirmed children.

**Supplementary Table 3:** Total differentially expressed genes in DN, Unstim DP, CD69^-^IFNγ^-^, CD69^+^IFNγ^-^ and CD69^+^IFNγ^+^ subsets.

**Supplementary Table 4:** Differences in the expression of cytokines/chemokines and their receptors in the HLADR^+^CD38^+^ CD8 T cell subsets

**Supplementary Table 5:** Differences in the expression of cytokines/chemokines and their receptors in the HLADR^+^CD38^+^ CD8 T cell subsets

**Supplementary Table 6:** Differences in the expression of genes related to cytotoxic effector functions in the HLADR^+^CD38^+^ CD8 T cell subsets

**Supplementary Table 7:** Differences in the expression of genes involved in TCR signaling / co-stimulation in the HLADR^+^CD38^+^ CD8 T cell subsets

**Supplementary Table 8:** Differences in the expression of genes involved in transcription, translation, metabolism, and proliferation in the CD69^+^IFNγ^+^ subset.

**Supplementary Table 9:** Characteristics of patients with DI, DW and SD.

## References

1. Bhatt S, Gething PW, Brady OJ, Messina JP, Farlow AW, Moyes CL, Drake JM, Brownstein JS, Hoen AG, Sankoh O, Myers MF, George DB, Jaenisch T, Wint GR, Simmons CP, Scott TW, Farrar JJ, Hay SI. 2013. The global distribution and burden of dengue. Nature 496:504–7.

2. Gupta E, Ballani N. 2014. Current perspectives on the spread of dengue in India. Infect Drug Resist 7:337–42.

3. WHO/TDR. 2009. Dengue: guidelines for diagnosis, treatment, prevention and control.

4. Halstead SB. 1979. In vivo enhancement of dengue virus infection in rhesus monkeys by passively transferred antibody. J Infect Dis 140:527–33.

5. Kliks SC, Nisalak A, Brandt WE, Wahl L, Burke DS. 1989. Antibody-dependent enhancement of dengue virus growth in human monocytes as a risk factor for dengue hemorrhagic fever. Am J Trop Med Hyg 40:444–51.

6. Mangada MM, Rothman AL. 2005. Altered cytokine responses of dengue-specific CD4+ T cells to heterologous serotypes. J Immunol 175:2676–83.

7. Mongkolsapaya J, Dejnirattisai W, Xu XN, Vasanawathana S, Tangthawornchaikul N, Chairunsri A, Sawasdivorn S, Duangchinda T, Dong T, Rowland-Jones S, Yenchitsomanus PT, McMichael A, Malasit P, Screaton G. 2003. Original antigenic sin and apoptosis in the pathogenesis of dengue hemorrhagic fever. Nat Med 9:921–7.

8. Pang T, Cardosa MJ, Guzman MG. 2007. Of cascades and perfect storms: the immunopathogenesis of dengue haemorrhagic fever-dengue shock syndrome (DHF/DSS). Immunol Cell Biol 85:43–5.

9. Sabin AB. 1952. Research on dengue during World War II. Am J Trop Med Hyg 1:30–50.

10. Snow GE, Haaland B, Ooi EE, Gubler DJ. 2014. Review article: Research on dengue during World War II revisited. Am J Trop Med Hyg 91:1203–17.

11. Wilder-Smith A. 2020. Dengue vaccine development: status and future. Bundesgesundheitsblatt Gesundheitsforschung Gesundheitsschutz 63:40–44.

12. Katzelnick LC, Gresh L, Halloran ME, Mercado JC, Kuan G, Gordon A, Balmaseda A, Harris E. 2017. Antibody-dependent enhancement of severe dengue disease in humans. Science 358:929–932.

13. Duangchinda T, Dejnirattisai W, Vasanawathana S, Limpitikul W, Tangthawornchaikul N, Malasit P, Mongkolsapaya J, Screaton G. 2010. Immunodominant T-cell responses to dengue virus NS3 are associated with DHF. Proc Natl Acad Sci U S A 107:16922–7.

14. Dung NT, Duyen HT, Thuy NT, Ngoc TV, Chau NV, Hien TT, Rowland-Jones SL, Dong T, Farrar J, Wills B, Simmons CP. 2010. Timing of CD8+ T cell responses in relation to commencement of capillary leakage in children with dengue. J Immunol 184:7281–7.

15. Elong Ngono A, Chen HW, Tang WW, Joo Y, King K, Weiskopf D, Sidney J, Sette A, Shresta S. 2016. Protective Role of Cross-Reactive CD8 T Cells Against Dengue Virus Infection. EBioMedicine 13:284–293.

16. Grifoni A, Voic H, Yu ED, Mateus J, Yan Fung KM, Wang A, Seumois G, De Silva AD, Tennekon R, Premawansa S, Premawansa G, Tippalagama R, Wijewickrama A, Chawla A, Greenbaum J, Peters B, Pandurangan V, Weiskopf D, Sette A. 2022. Transcriptomics of Acute DENV-Specific CD8+ T Cells Does Not Support Qualitative Differences as Drivers of Disease Severity. Vaccines (Basel) 10.

17. Lam JH, Chua YL, Lee PX, Martinez Gomez JM, Ooi EE, Alonso S. 2017. Dengue vaccine-induced CD8+ T cell immunity confers protection in the context of enhancing, interfering maternal antibodies. JCI Insight 2.

18. Weiskopf D, Angelo MA, de Azeredo EL, Sidney J, Greenbaum JA, Fernando AN, Broadwater A, Kolla RV, De Silva AD, de Silva AM, Mattia KA, Doranz BJ, Grey HM, Shresta S, Peters B, Sette A. 2013. Comprehensive analysis of dengue virus-specific responses supports an HLA-linked protective role for CD8+ T cells. Proc Natl Acad Sci U S A 110:E2046–53.

19. Chandele A, Sewatanon J, Gunisetty S, Singla M, Onlamoon N, Akondy RS, Kissick HT, Nayak K, Reddy ES, Kalam H, Kumar D, Verma A, Panda H, Wang S, Angkasekwinai N, Pattanapanyasat K, Chokephaibulkit K, Medigeshi GR, Lodha R, Kabra S, Ahmed R, Murali-Krishna K. 2016. Characterization of Human CD8 T Cell Responses in Dengue Virus-Infected Patients from India. J Virol 90:11259–11278.

20. de Matos AM, Carvalho KI, Rosa DS, Villas-Boas LS, da Silva WC, Rodrigues CL, Oliveira OM, Levi JE, Araujo ES, Pannuti CS, Luna EJ, Kallas EG. 2015. CD8+ T lymphocyte expansion, proliferation and activation in dengue fever. PLoS Negl Trop Dis 9:e0003520.

21. Tian Y, Grifoni A, Sette A, Weiskopf D. 2019. Human T Cell Response to Dengue Virus Infection. Front Immunol 10:2125.

22. Akondy RS, Monson ND, Miller JD, Edupuganti S, Teuwen D, Wu H, Quyyumi F, Garg S, Altman JD, Del Rio C, Keyserling HL, Ploss A, Rice CM, Orenstein WA, Mulligan MJ, Ahmed R. 2009. The yellow fever virus vaccine induces a broad and polyfunctional human memory CD8+ T cell response. J Immunol 183:7919–30.

23. Waickman AT, Victor K, Li T, Hatch K, Rutvisuttinunt W, Medin C, Gabriel B, Jarman RG, Friberg H, Currier JR. 2019. Dissecting the heterogeneity of DENV vaccine-elicited cellular immunity using single-cell RNA sequencing and metabolic profiling. Nat Commun 10:3666.

24. Matsuo K, Kitahata K, Kawabata F, Kamei M, Hara Y, Takamura S, Oiso N, Kawada A, Yoshie O, Nakayama T. 2018. A Highly Active Form of XCL1/Lymphotactin Functions as an Effective Adjuvant to Recruit Cross-Presenting Dendritic Cells for Induction of Effector and Memory CD8(+) T Cells. Front Immunol 9:2775.

25. Wang M, Windgassen D, Papoutsakis ET. 2008. Comparative analysis of transcriptional profiling of CD3+, CD4+ and CD8+ T cells identifies novel immune response players in T-cell activation. BMC Genomics 9:225.

26. Mehraj V, Ramendra R, Isnard S, Dupuy FP, Lebouche B, Costiniuk C, Thomas R, Szabo J, Baril JG, Trottier B, Cote P, LeBlanc R, Durand M, Chartrand-Lefebvre C, Kema I, Zhang Y, Finkelman M, Tremblay C, Routy JP. 2019. CXCL13 as a Biomarker of Immune Activation During Early and Chronic HIV Infection. Front Immunol 10:289.

27. Ogasawara H, Takeda-Hirokawa N, Sekigawa I, Hashimoto H, Kaneko Y, Hirose S. 1999. Inhibitory effect of interleukin-16 on interleukin-2 production by CD4+ T cells. Immunology 96:215–9.

28. Bie Q, Jin C, Zhang B, Dong H. 2017. IL-17B: A new area of study in the IL-17 family. Mol Immunol 90:50–56.

29. Anuradha R, Munisankar S, Dolla C, Kumaran P, Nutman TB, Babu S. 2016. Modulation of CD4+ and CD8+ T-Cell Function by Interleukin 19 and Interleukin 24 During Filarial Infections. J Infect Dis 213:811–5.

30. Bezie S, Picarda E, Ossart J, Tesson L, Usal C, Renaudin K, Anegon I, Guillonneau C. 2015. IL-34 is a Treg-specific cytokine and mediates transplant tolerance. J Clin Invest 125:3952–64.

31. Cavalli G, Tengesdal IW, Gresnigt M, Nemkov T, Arts RJW, Dominguez-Andres J, Molteni R, Stefanoni D, Cantoni E, Cassina L, Giugliano S, Schraa K, Mills TS, Pietras EM, Eisenmensser EZ, Dagna L, Boletta A, D’Alessandro A, Joosten LAB, Netea MG, Dinarello CA. 2021. The anti-inflammatory cytokine interleukin-37 is an inhibitor of trained immunity. Cell Rep 35:108955.

32. Yuan ZC, Xu WD, Liu XY, Liu XY, Huang AF, Su LC. 2019. Biology of IL-36 Signaling and Its Role in Systemic Inflammatory Diseases. Front Immunol 10:2532.

33. Daynes RA, Dowell T, Araneo BA. 1991. Platelet-derived growth factor is a potent biologic response modifier of T cells. J Exp Med 174:1323–33.

34. Lyons RM, Moses HL. 1990. Transforming growth factors and the regulation of cell proliferation. Eur J Biochem 187:467–73.

35. Wei F, Zhang Y, Zhao W, Yu X, Liu CJ. 2014. Progranulin facilitates conversion and function of regulatory T cells under inflammatory conditions. PLoS One 9:e112110.

36. Tominaga K, Yoshimoto T, Torigoe K, Kurimoto M, Matsui K, Hada T, Okamura H, Nakanishi K. 2000. IL-12 synergizes with IL-18 or IL-1beta for IFN-gamma production from human T cells. Int Immunol 12:151–60.

37. Kurata R, Shimizu K, Cui X, Harada M, Isagawa T, Semba H, Ishihara J, Yamada K, Nagai J, Yoshida Y, Takeda N, Maemura K, Yonezawa T. 2020. Novel Reporter System Monitoring IL-18 Specific Signaling Can Be Applied to High-Throughput Screening. Mar Drugs 18.

38. Groom JR, Luster AD. 2011. CXCR3 in T cell function. Exp Cell Res 317:620–31.

39. Rennier K, Shin WJ, Krug E, Virdi G, Pachynski RK. 2020. Chemerin Reactivates PTEN and Suppresses PD-L1 in Tumor Cells via Modulation of a Novel CMKLR1-mediated Signaling Cascade. Clin Cancer Res 26:5019–5035.

40. Robinson RT. 2015. IL12Rbeta1: the cytokine receptor that we used to know. Cytokine 71:348–59.

41. Lee J, Aoki T, Thumkeo D, Siriwach R, Yao C, Narumiya S. 2019. T cell-intrinsic prostaglandin E2-EP2/EP4 signaling is critical in pathogenic TH17 cell-driven inflammation. J Allergy Clin Immunol 143:631–643.

42. Zhu N, Yang Y, Wang H, Tang P, Zhang H, Sun H, Gong L, Yu Z. 2022. CSF2RB Is a Unique Biomarker and Correlated With Immune Infiltrates in Lung Adenocarcinoma. Front Oncol 12:822849.

43. Gusscott S, Jenkins CE, Lam SH, Giambra V, Pollak M, Weng AP. 2016. IGF1R Derived PI3K/AKT Signaling Maintains Growth in a Subset of Human T-Cell Acute Lymphoblastic Leukemias. PLoS One 11:e0161158.

44. Fuchs YF, Sharma V, Eugster A, Kraus G, Morgenstern R, Dahl A, Reinhardt S, Petzold A, Lindner A, Lobel D, Bonifacio E. 2019. Gene Expression-Based Identification of Antigen-Responsive CD8(+) T Cells on a Single-Cell Level. Front Immunol 10:2568.

45. Sohar I, Sleat D, Gong Liu C, Ludwig T, Lobel P. 1998. Mouse mutants lacking the cation-independent mannose 6-phosphate/insulin-like growth factor II receptor are impaired in lysosomal enzyme transport: comparison of cation-independent and cation-dependent mannose 6-phosphate receptor-deficient mice. Biochem J 330 (Pt 2):903–8.

46. Croft M. 2009. The role of TNF superfamily members in T-cell function and diseases. Nat Rev Immunol 9:271–85.

47. Wikenheiser DJ, Stumhofer JS. 2016. ICOS Co-Stimulation: Friend or Foe? Front Immunol 7:304.

48. Wang Z, Yin N, Zhang Z, Zhang Y, Zhang G, Chen W. 2017. Upregulation of T-cell Immunoglobulin and Mucin-Domain Containing-3 (Tim-3) in Monocytes/Macrophages Associates with Gastric Cancer Progression. Immunol Invest 46:134–148.

49. Blackburn SD, Shin H, Haining WN, Zou T, Workman CJ, Polley A, Betts MR, Freeman GJ, Vignali DA, Wherry EJ. 2009. Coregulation of CD8+ T cell exhaustion by multiple inhibitory receptors during chronic viral infection. Nat Immunol 10:29–37.

50. Zhang X, Yu Y, Bai B, Wang T, Zhao J, Zhang N, Zhao Y, Wang X, Wang B. 2020. PTPN22 interacts with EB1 to regulate T-cell receptor signaling. FASEB J 34:8959–8974.

51. Ferris RL, Lu B, Kane LP. 2014. Too much of a good thing? Tim-3 and TCR signaling in T cell exhaustion. J Immunol 193:1525–30.

52. Sun F, Yue TT, Yang CL, Wang FX, Luo JH, Rong SJ, Zhang M, Guo Y, Xiong F, Wang CY. 2021. The MAPK dual specific phosphatase (DUSP) proteins: A versatile wrestler in T cell functionality. Int Immunopharmacol 98:107906.

53. Wan X, Zhang J, Luo H, Shi G, Kapnik E, Kim S, Kanakaraj P, Wu J. 2002. A TNF family member LIGHT transduces costimulatory signals into human T cells. J Immunol 169:6813–21.

54. Liu X, Zhelev D, Adams C, Chen C, Mellors JW, Dimitrov DS. 2021. Effective killing of cells expressing CD276 (B7-H3) by a bispecific T cell engager based on a new fully human antibody. Transl Oncol 14:101232.

55. Rollins MR, Gibbons Johnson RM. 2017. CD80 Expressed by CD8(+) T Cells Contributes to PD-L1-Induced Apoptosis of Activated CD8(+) T Cells. J Immunol Res 2017:7659462.

56. Barazeghi E, Hellman P, Westin G, Stalberg P. 2019. PTPRM, a candidate tumor suppressor gene in small intestinal neuroendocrine tumors. Endocr Connect 8:1126–1135.

57. Stanford SM, Rapini N, Bottini N. 2012. Regulation of TCR signalling by tyrosine phosphatases: from immune homeostasis to autoimmunity. Immunology 137:1–19.

58. Kowalczyk A, Kleniewska P, Kolodziejczyk M, Skibska B, Goraca A. 2015. The role of endothelin-1 and endothelin receptor antagonists in inflammatory response and sepsis. Arch Immunol Ther Exp (Warsz) 63:41–52.

59. Son HJ, An CH, Yoo NJ, Lee SH. 2020. Tight Junction-Related CLDN5 and CLDN6 Genes, and Gap Junction-Related GJB6 and GJB7 Genes Are Somatically Mutated in Gastric and Colorectal Cancers. Pathol Oncol Res 26:1983–1987.

60. Martin E, Palmic N, Sanquer S, Lenoir C, Hauck F, Mongellaz C, Fabrega S, Nitschke P, Esposti MD, Schwartzentruber J, Taylor N, Majewski J, Jabado N, Wynn RF, Picard C, Fischer A, Arkwright PD, Latour S. 2014. CTP synthase 1 deficiency in humans reveals its central role in lymphocyte proliferation. Nature 510:288–92.

61. Bachmann AS, Geerts D. 2018. Polyamine synthesis as a target of MYC oncogenes. J Biol Chem 293:18757–18769.

62. Goswami MT, Chen G, Chakravarthi BV, Pathi SS, Anand SK, Carskadon SL, Giordano TJ, Chinnaiyan AM, Thomas DG, Palanisamy N, Beer DG, Varambally S. 2015. Role and regulation of coordinately expressed de novo purine biosynthetic enzymes PPAT and PAICS in lung cancer. Oncotarget 6:23445–61.

63. Aggarwal C, Saini K, Reddy ES, Singla M, Nayak K, Chawla YM, Maheshwari D, Singh P, Sharma P, Bhatnagar P, Kumar S, Gottimukkala K, Panda H, Gunisetty S, Davis CW, Kissick HT, Kabra SK, Lodha R, Medigeshi GR, Ahmed R, Murali-Krishna K, Chandele A. 2021. Immunophenotyping and Transcriptional Profiling of Human Plasmablasts in Dengue. J Virol 95:e0061021.

64. Arreola R, Valderrama B, Morante ML, Horjales E. 2003. Two mammalian glucosamine-6-phosphate deaminases: a structural and genetic study. FEBS Lett 551:63–70.

65. Hickman TL, Choi E, Whiteman KR, Muralidharan S, Pai T, Johnson T, Parikh A, Friedman T, Gilbert M, Shen B, Barron L, McGinness KE, Ettenberg SA, Motz GT, Weiss GJ, Jensen-Smith A. 2022. BOXR1030, an anti-GPC3 CAR with exogenous GOT2 expression, shows enhanced T cell metabolism and improved anti-cell line derived tumor xenograft activity. PLoS One 17:e0266980.

66. Hou T, Zhang R, Jian C, Ding W, Wang Y, Ling S, Ma Q, Hu X, Cheng H, Wang X. 2019. NDUFAB1 confers cardio-protection by enhancing mitochondrial bioenergetics through coordination of respiratory complex and supercomplex assembly. Cell Res 29:754–766.

67. Cannarile MA, Lind NA, Rivera R, Sheridan AD, Camfield KA, Wu BB, Cheung KP, Ding Z, Goldrath AW. 2006. Transcriptional regulator Id2 mediates CD8+ T cell immunity. Nat Immunol 7:1317–25.

68. Luo CT, Osmanbeyoglu HU, Do MH, Bivona MR, Toure A, Kang D, Xie Y, Leslie CS, Li MO. 2017. Ets transcription factor GABP controls T cell homeostasis and immunity. Nat Commun 8:1062.

69. Lake RJ, Tsai PF, Choi I, Won KJ, Fan HY. 2014. RBPJ, the major transcriptional effector of Notch signaling, remains associated with chromatin throughout mitosis, suggesting a role in mitotic bookmarking. PLoS Genet 10:e1004204.

70. Johnston J, Basatvat S, Ilyas Z, Francis S, Kiss-Toth E. 2015. Tribbles in inflammation. Biochem Soc Trans 43:1069–74.

71. Lu G, Tian S, Sun Y, Dong J, Wang N, Zeng J, Nie Y, Wu K, Han Y, Feng B, Shang Y. 2021. NEK9, a novel effector of IL-6/STAT3, regulates metastasis of gastric cancer by targeting ARHGEF2 phosphorylation. Theranostics 11:2460–2474.

72. Choi H, Cho SY, Schwartz RH, Choi K. 2006. Dual effects of Sprouty1 on TCR signaling depending on the differentiation state of the T cell. J Immunol 176:6034–45.

73. Yigit B, Wang N, Ten Hacken E, Chen SS, Bhan AK, Suarez-Fueyo A, Katsuyama E, Tsokos GC, Chiorazzi N, Wu CJ, Burger JA, Herzog RW, Engel P, Terhorst C. 2019. SLAMF6 as a Regulator of Exhausted CD8(+) T Cells in Cancer. Cancer Immunol Res 7:1485–1496.

74. Dominguez CX, Amezquita RA, Guan T, Marshall HD, Joshi NS, Kleinstein SH, Kaech SM. 2015. The transcription factors ZEB2 and T-bet cooperate to program cytotoxic T cell terminal differentiation in response to LCMV viral infection. J Exp Med 212:2041–56.

75. Joshi NS, Cui W, Chandele A, Lee HK, Urso DR, Hagman J, Gapin L, Kaech SM. 2007. Inflammation directs memory precursor and short-lived effector CD8(+) T cell fates via the graded expression of T-bet transcription factor. Immunity 27:281–95.

76. Ahmed R, Akondy RS. 2011. Insights into human CD8(+) T-cell memory using the yellow fever and smallpox vaccines. Immunol Cell Biol 89:340–5.

77. Lechner F, Wong DK, Dunbar PR, Chapman R, Chung RT, Dohrenwend P, Robbins G, Phillips R, Klenerman P, Walker BD. 2000. Analysis of successful immune responses in persons infected with hepatitis C virus. J Exp Med 191:1499–512.

78. Welsh RM. 2001. Assessing CD8 T cell number and dysfunction in the presence of antigen. J Exp Med 193:F19–22.

79. Kaech SM, Hemby S, Kersh E, Ahmed R. 2002. Molecular and functional profiling of memory CD8 T cell differentiation. Cell 111:837–51.

80. Kaech SM, Tan JT, Wherry EJ, Konieczny BT, Surh CD, Ahmed R. 2003. Selective expression of the interleukin 7 receptor identifies effector CD8 T cells that give rise to long-lived memory cells. Nat Immunol 4:1191–8.

81. Kaech SM, Wherry EJ. 2007. Heterogeneity and cell-fate decisions in effector and memory CD8+ T cell differentiation during viral infection. Immunity 27:393–405.

82. Chng MHY, Lim MQ, Rouers A, Becht E, Lee B, MacAry PA, Lye DC, Leo YS, Chen J, Fink K, Rivino L, Newell EW. 2019. Large-Scale HLA Tetramer Tracking of T Cells during Dengue Infection Reveals Broad Acute Activation and Differentiation into Two Memory Cell Fates. Immunity 51:1119–1135 e5.

83. Chen H, Smith M, Herz J, Li T, Hasley R, Le Saout C, Zhu Z, Cheng J, Gronda A, Martina JA, Irusta PM, Karpova T, McGavern DB, Catalfamo M. 2021. The role of protease-activated receptor 1 signaling in CD8 T cell effector functions. iScience 24:103387.

